# Developmental series of gene expression clarifies maternal mRNA provisioning and maternal-to-zygotic transition in the reef-building coral *Montipora capitata*

**DOI:** 10.1101/2021.04.14.439692

**Authors:** E Chille, E Strand, M Neder, V Schmidt, M Sherman, T Mass, HM Putnam

## Abstract

**Background:** Maternal mRNA provisioning of oocytes regulates early embryogenesis. Maternal transcripts are degraded as zygotic genome activation (ZGA) intensifies, a phenomenon known as the maternal-to-zygotic transition (MZT). Here, we examine gene expression over nine developmental stages in the Pacific rice coral, *Montipora capitata*, from eggs and embryos at 1, 4, 9, 14, 22, and 36 hours-post-fertilization (hpf), as well as swimming larvae (9d), and adult colonies.

**Results:** Weighted Gene Coexpression Network Analysis revealed four expression peaks, identifying the maternal complement, two waves of the MZT, and adult expression. Gene ontology enrichment revealed maternal mRNAs are dominated by cell division, methylation, biosynthesis, metabolism, and protein/RNA processing and transport functions. The first MZT wave occurs from ∼4-14 hpf and is enriched in terms related to biosynthesis, methylation, cell division, and transcription. In contrast, functional enrichment in the second MZT wave, or ZGA, from 22 hpf-9dpf, includes ion/peptide transport and cell signaling. Finally, adult expression is enriched for functions related to signaling, metabolism, and ion/peptide transport. Our proposed MZT timing is further supported by expression of enzymes involved in zygotic transcriptional repression (Kaiso) and activation (Sox2), which peak at 14 hpf and 22 hpf, respectively. Further, DNA methylation writing (DNMT3a) and removing enzymes (TET1) peak and remain stable past ∼4 hpf, indicating that methylome programming occurs before 4 hpf.

**Conclusions:** Our high-resolution insight into the coral maternal mRNA and MZT provides essential information regarding setting the stage for, and the sensitivity of, developmental success and parental carryover effects under increasing environmental stress.

## Background

Under escalating threats from anthropogenic climate change, coral reefs are experiencing massive population declines globally [1]. Contributing to the global destabilization of coral reef ecosystems is the decreased fitness of coral during its early development [2–5] due to anthropogenic climate change [6–8]. The majority of corals (∼85%) are broadcast spawners [9], releasing eggs and sperm into the water column where fertilization and embryogenesis occur [10]. Over the course of hours to days, embryos develop into planula larvae [10] and may spend several hours to weeks in the water column, or searching the benthos for an appropriate location to settle and metamorphose into coral spat [11]. This prolonged pelagic period constitutes an ontogenetic bottleneck for corals and other broadcast-spawning marine organisms, during which they are particularly sensitive to the climate-related stressors [6,7].

Acute temperature stress can be lethal to coral embryos and larvae, while ocean acidification can have sub-lethal effects [7], including decreased fertilization efficiency [12,13] and metabolic depression [14–17]. Other conditions co-occurring with climate change, including high and low salinity, hypoxia, and exposure to UV radiation, can also decrease fitness during development [7,8]. The cumulative effects of these unprecedented stressors on early development may compromise recruitment, the process in which offspring repopulate and replenish declining populations, further contributing to the destabilization of coral reef ecosystems [2].

Several studies on the sensitivity of embryonic development have shown that the earliest life stages are robust to climate-related stressors, but embryos become more vulnerable as gastrula and planula [6,7,18–22]. This has generally been attributed to the loss of maternal defenses, such as heat shock proteins, during the course of development [6,7,18–22]. During oogenesis, maternally-derived mRNAs (or parentally-derived in the case of hermaphroditic species), along with other gene products (i.e. transcription factors, cofactors), are loaded into eggs [23,24]. As gametes and early embryos do not yet contain the molecular machinery necessary to regulate gene expression, these mRNAs direct early development until the zygotic molecular machinery is able to take over [23]. The maternal mRNA complement is the result of selection in response to life history [24,25], and can be primed to support embryonic resilience to predictable stressors [26–28]. For example, it has been shown that in cichlids, the maternal mRNA complement differs by trophic specialization [25]. Additionally, many organisms can prime the immune response of their offspring by providing increased levels of mRNAs coding for immune response-related enzymes [25,27–31]. In terms of marine invertebrates, the annelid *Hediste diversicolor*, was found to store higher levels of the antimicrobial peptide hedistin in the oocytes after an immune challenge [28], and sea urchins with different life history strategies (lecithotrophy or planktotrophy) were found to differentially provision oocytes with mRNAs related to oxidative stress [32].

Maternal provisioning may explain the resilience of early coral embryos to expected stressors, and is a mechanism of coral reef resilience that is only starting to be fully investigated [33–36]. However, there is growing concern that maternal provisioning may not provide adequate defense against rapid climate change, especially for organisms living at the edge of their physiological limits [7,22,24,37]. For example, coral bleaching from temperature stress can negatively impact maternal provisioning of protein, lipid, mycosporine-like amino acids and carotenoids in the eggs [38]. Consequently, characterization of the coral maternal mRNA complement can additionally provide essential information regarding setting the stage for, and the sensitivity of, developmental success under increasing environmental stressors.

While maternal provisioning may protect the embryo from expected stressors early on, these gene products only persist for a limited time, after which point embryo vulnerability appears to intensify [6,7,18–22]. In a process known as the maternal-to-zygotic transition (MZT), maternal gene products are degraded as zygotic transcription activates and intensifies [39–41]. The loss of maternal defenses during this process has been posited to contribute to higher morbidity and mortality in fish gastrula, sea urchin blastula/gastrula, as well as many marine invertebrate larvae during exposure to environmental stressors [7,20–22,37].

The MZT and many of the molecular mechanisms underlying its regulation are conserved in metazoans [39,40,42–44]. Maternal mRNA clearance is mediated both maternally, primarily by the enzyme Smaug [45,46], and zygotically [40]. Transcriptional repressors, including the maternally-derived enzyme Kaiso [47,48], prevent zygotic transcription until a stable, open chromatin state is achieved [49]. Once this occurs, ZGA begins gradually and ramps in intensity as it progresses [39,40]. The low-intensity beginning of ZGA (the “first” or “minor” wave) is controlled by maternal transcription factors, including Sox2 [50], while the ramping of ZGA to higher intensities (the “major wave”) is triggered by the clearance of the majority of maternal transcripts and the lengthening of the cell cycle through the degradation of cell-cycle regulators such as Cyclin-B [39,51]. Among the first of the enzymes shown to be zygotically expressed in sea urchins and other organisms are those involved in bauplan formation, including Wnt8 and Brachyury (TBXT) [52–54]. Upon culmination of the MZT, zygotically-derived gene products regulate developmental progression and homeostasis [39]. Characterizing the timeline of the MZT in reef-building corals in regards to these conserved mechanisms will help to establish the time period during which maternal defenses are active, and at which point coral embryos may become more susceptible to environmental perturbations.

Enzymes linked to epigenetic modification are also shown to be key players in the repression and activation of zygotic transcription, with increases in DNA methylation and simultaneous decreases in chromatin accessibility associated with the onset of ZGA [40,55]. Additionally, these enzymes confer potential mechanisms for cross-generational plasticity, which may additionally buffer reef-building corals and other organisms from the effects of unprecedented stressors [56,57]. Several key epigenetic modification associated enzymes that provide transcriptional regulation capacity include: DNA methyl-transferases (DNMT3a, DNMT1), the histone deacetylase recruiting enzyme ubiquitin-like containing PHD and RING finger domains 1 (UHRF1), and methyl-binding domains (MBD2, MBD3), as well as the active demethylation enzyme ten-eleven translocation enzyme (TET1) and chromatin regulation enzyme brahma-related gene-1 (BRG1). DNMTs are responsible for laying down DNA methylation *de novo* (DNMT3a) and maintaining DNA methylation through mitosis (DNMT1) [58], while TET1 actively removes DNA methylation [59]. Depletion of DNMT1 can lead to the premature onset of ZGA in *Xenopus* embryos [60]. The protein UHRF1 is associated with DNMT1, marking hemimethylated DNA for DNMT1 activity [61]. The closely-related MBD2 and MBD3 proteins are also involved in transcriptional repression as components of the Mi-2/NuRD corepressor complex [62–64]. In mice, it has been suggested that MBD3 is essential for embryo viability as a key component of the Mi-2/NuRD corepressor complex, while MBD2 may be a functionally redundant cofactor [64]. Finally, BRG1 proteins are derived from the maternal mRNA complement and are suggested to be the first gene required for ZGA in mammals [65]. BRG1 is generally associated with increasing chromatin accessibility because of its role in breaking the bonds between DNA and histones through ATP hydrolysis. Analyzing the expression profiles of these specific enzymes involved in epigenetic modification may provide further support on the epigenetic state necessary for ZGA.

Here we used the knowledge on the well-documented spawning of the Pacific rice coral *Montipora capitata* in Hawai□i [66,67] and its physical development [68] to examine developmental timing through the lens of gene expression. In this study, we examine gene expression dynamics across nine life stages from pre-fertilization to adult, focusing on shifts in the gene expression network within the first 24 hours post-spawning. In doing so, we aim to characterize the *M. capitata* maternal mRNA complement, as well as the timing and functional patterns of ZGA and the MZT. This high resolution analysis of the *M. capitata* developmental transcriptome will provide a basis from which to investigate how its early developmental stages respond to environmental stressors, as well as the potential for carryover or latent effects due to parental history.

## Results

### RNA sequencing, quality control, and mapping

TruSeq Illumina Stranded Paired-End (PE) sequencing of cDNA libraries prepared following polyA enrichment resulted in 935,000,000 PE reads with an average of 19,392,156 reads per sample. After quality filtering and adapter removal, 694,800,000 PE reads remained with an average of 14,241,176 reads per sample. The average alignment rate was 79.28 ± 1.945384% mean ± sem. The GFFcompare results showed that a total of 63,227 genes queried from the mapped GTF files were matched to the 63,227 genes in the reference genome [69]. Altogether, there were 40,883 matching intron chains, 63,218 matching loci, and zero missing elements.

### Global gene expression patterns

Pre-filtering to retain genes with counts over 10 in at least 8.3% of samples (i.e., the proportion representing a single life stage) resulted in 32,772 genes for statistical analysis. A principal components analysis conducted with this gene set shows significant differences in global gene expression between time points with the exception of samples corresponding to unfertilized and fertilized eggs, which cluster together. Differences in gene expression between life stages is primarily explained by PC1 and PC2, which account for 71% and 14% of variation in expression, respectively (Fig 2). WGCNA assigned genes into 34 modules, with 48 to 11,142 genes in each module, one of which, module Grey, contains 29 genes that did not fit with any other group. Modules exhibited nine distinct expression profiles (Fig 3 left axis) based on module clustering by eigengene dissimilarity. Further, clustering of life stages by module-trait correlation identified four expression groups, which we call developmental clusters throughout this manuscript (Fig 3, clustering on top axis). The first developmental cluster, the “Maternal” modules, contains unfertilized and fertilized eggs (modules grey, coral1, lightslateblue, mediumpurple3, antiquewhite2, antiquewhite4, thistle, honeydew1, and midnightblue). The second developmental cluster, the “ZGA Wave 1” modules, contains the life stages cleavage, prawn chip, and early gastrula (modules magenta4, indianred3, blue2, plum3, blue4, skyblue1, brown2, coral, darkslateblue, plum4, and violet). The third developmental cluster, the “ZGA Wave 2” modules, contains the life stages mid-gastrula, late gastrula, and planula (modules magenta4, skyblue1, darkseagreen, darkslateblue, thistle4, salmon4, mediumpurple1, sienna3, salmon, and blue). Finally, the fourth developmental cluster, “Adult”, contains only the adult samples (modules antiquewhite4, thistle, navajowhite1, blue, cyan, blueviolet, and ivory).

**Figure 1.**
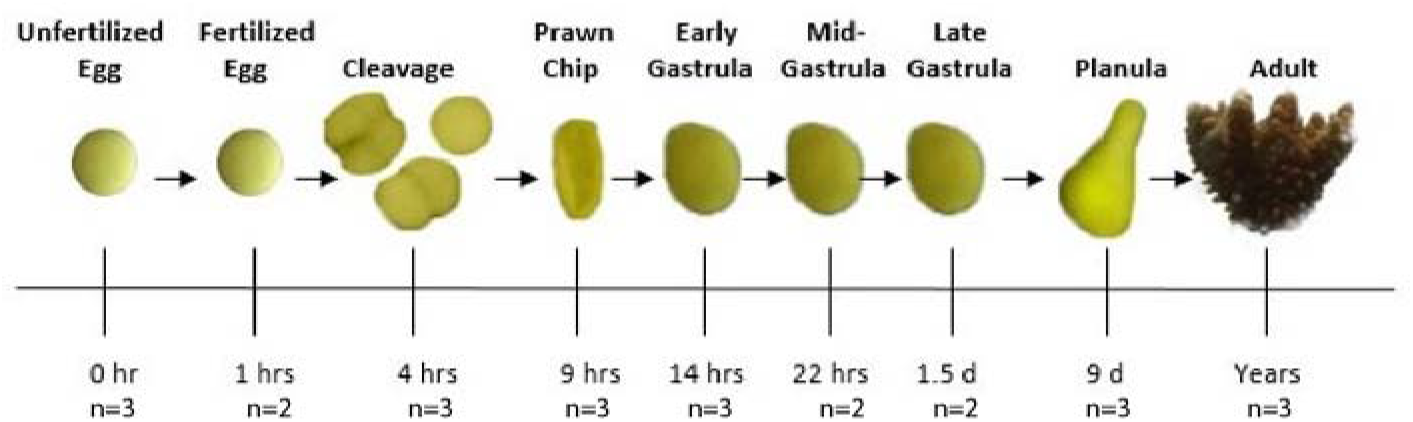
Timeline of sample collection including example photographs of life each stage.

**Figure 2.**
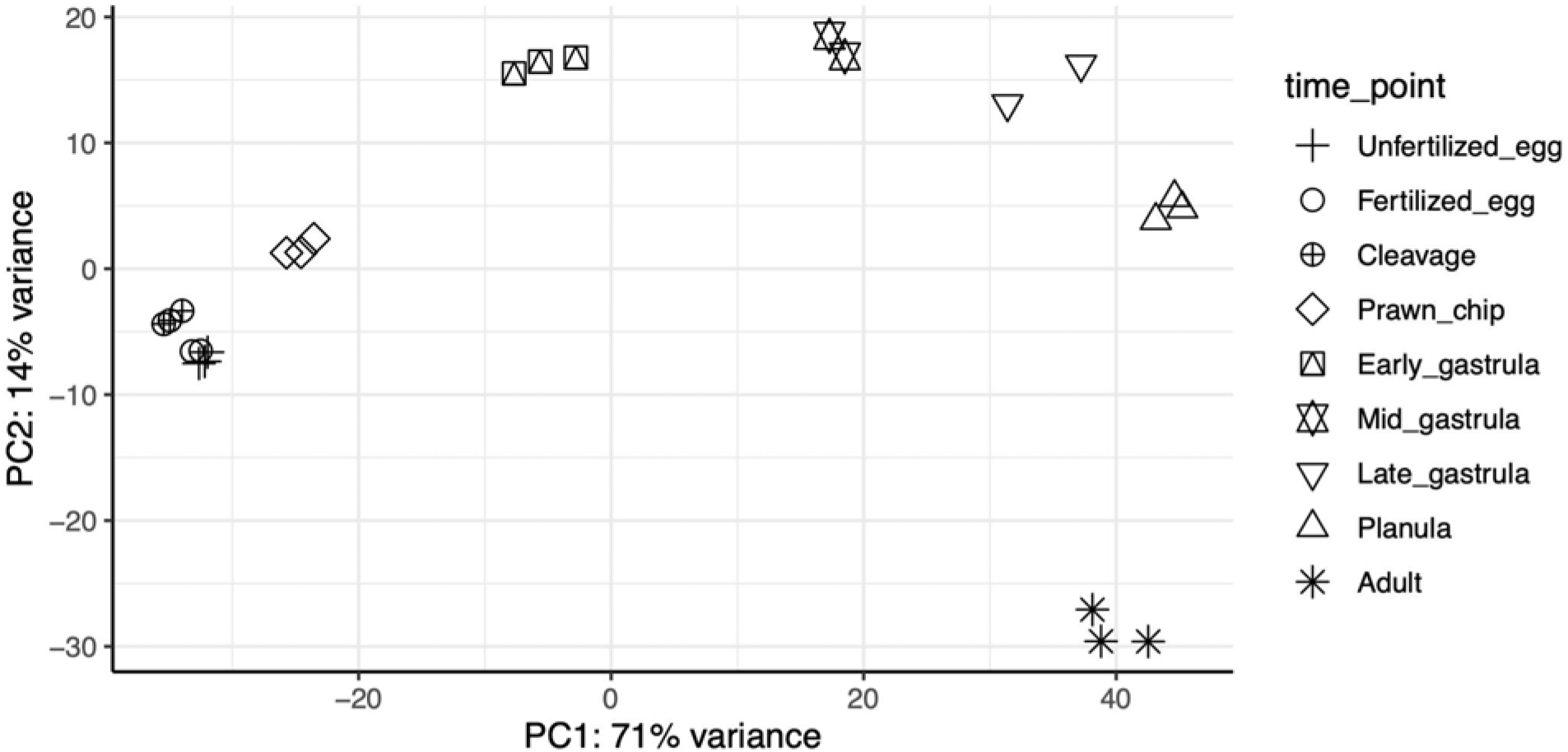
Principal coordinates analysis based on sample-to-sample distance computed from all genes passing a low counts filter, wherein a gene must have a count of 10 or greater in at least 2 out of the 24 samples (pOverA 0.083, 10).

**Figure 3.**
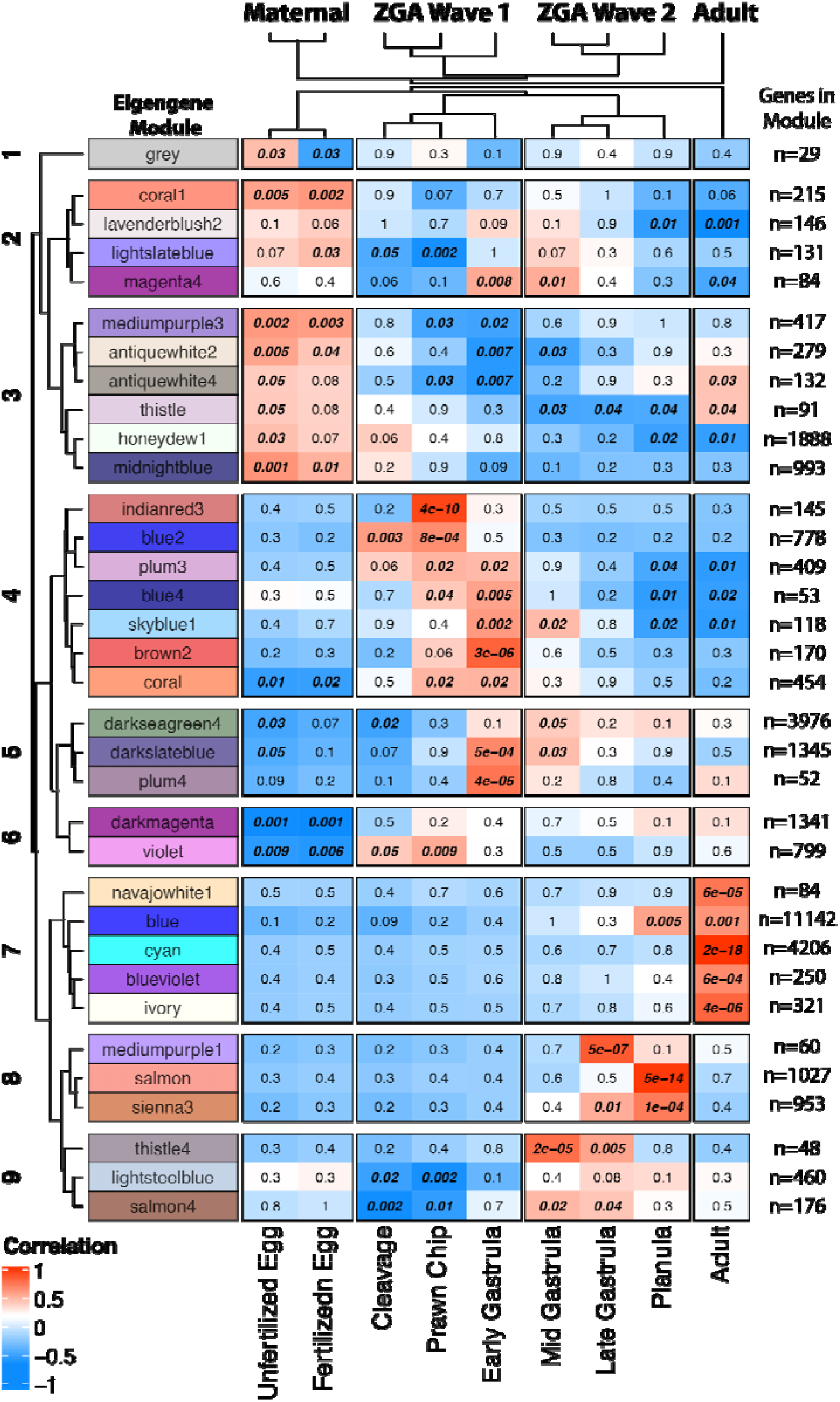
Clustered heatmap of WGCNA module-trait correlations ranging from correlation values of −1 (dark blue) to 0 (white) to +1 (red). Developmental expression clusters are shown on top, while sampled life stages are indicated on bottom. Module clusters 1-9 were identified via a distance matrix computed with hclust using module eigengenes.

### Functional annotation

To provide up-to-date gene ontology (GO) annotation for functional enrichment analysis, functional annotation was undertaken using DIAMOND, InterProScan, Blast2GO, and Uniprot [70–73]. Sequence alignment using DIAMOND resulted in 55,217 significant pairwise alignments for the total 63,227 sequences queried, with a median e-value of 3.100000e-72 and a median bitscore of 282.3. Blast2GO mapped 4,205 sequences to one or more GO terms. Additionally, a UniProt search of the protein identifiers obtained through DIAMOND matched 2,351 sequences with one or more GO terms (7,408 total terms). Finally, annotation with InterProScan matched 49,338 out of the 63,227 query sequences to entries in the InterProScan databases, 20,603 of which were associated with one or more GO terms (47,726 total terms). In total, 3,264 unique GO terms were mapped to 23,107 genes, with a total of 55,974 annotations to the *M. capitata* transcriptome (Table S1).

### Functional Enrichment

Functional enrichment analysis and subsequent semantic similarity analysis of biological process and molecular function terms was undertaken for each life stage (Table S2) and for the four developmental clusters: Maternal, ZGA Wave 1, ZGA Wave 2, and Adult (Fig. 4, Fig. 5). The maternal complement has 92 enriched biological processes (Table S3) primarily related to cell division, DNA repair, methylation, biosynthesis, metabolism, and protein/RNA processing and transport and 92 enriched molecular functions (Table S3) primarily related to binding, translation, transmembrane transport, and enzymatic activity. The first wave of the ZGA is enriched in 55 biological process terms (Table S4) and 60 molecular function terms (Table S4). Biosynthesis activity remains high during the first wave of the ZGA, which is additionally associated with enrichment of GO terms related to RNA processing and histone assembly. The second wave of the ZGA is enriched in 22 biological process terms (Table S5) of transmembrane transport and DNA metabolic process, as well as 47 molecular function terms, corresponding to transport, oxidation-reduction process, and binding. Finally, the adult transcriptome is enriched in 31 biological process terms (Table S6) and 54 molecular functions. GO terms related to metabolism and transmembrane transport remain high in the adult transcriptome, in addition to terms related to cell signaling.

**Figure 4.**
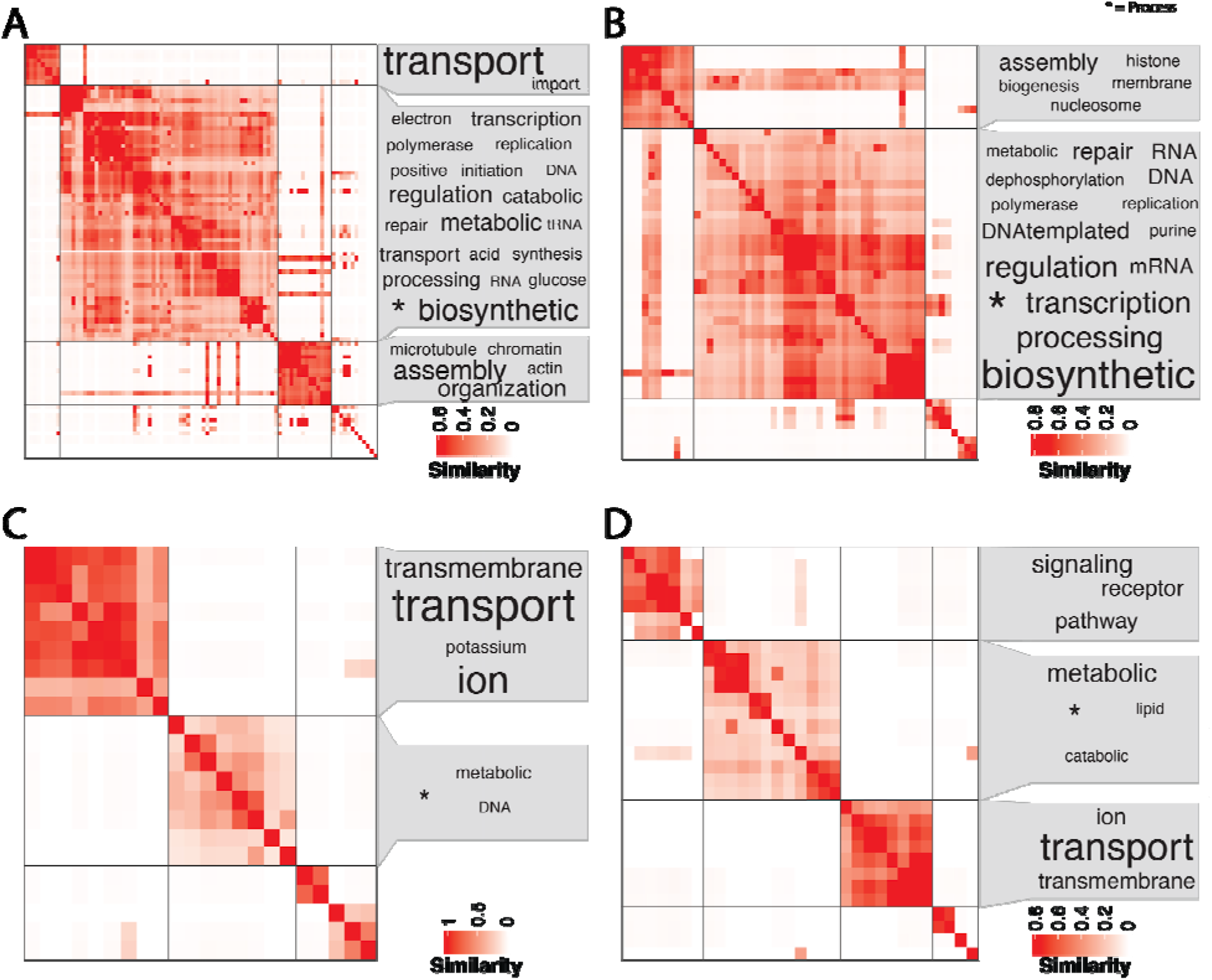
Clustered (binary cut) semantic similarity matrix of Biological Process gene ontology terms for all genes in the 9 modules identified as the A) maternal mRNA complement (n=92 terms), the 11 modules of the B) first wave of the MZT (n=55 terms), and the 10 modules of the C) second wave of the MZT transition (n=22 terms) and the 7 modules of D)“adult” expression (n=31 terms). Terms representing each cluster are shown in a word cloud, with size of term representing frequency of occurrence in the terms. The generic term, “Process” (identified by * in each graphic), was put as a footnote for visual clarity.

**Figure 5.**
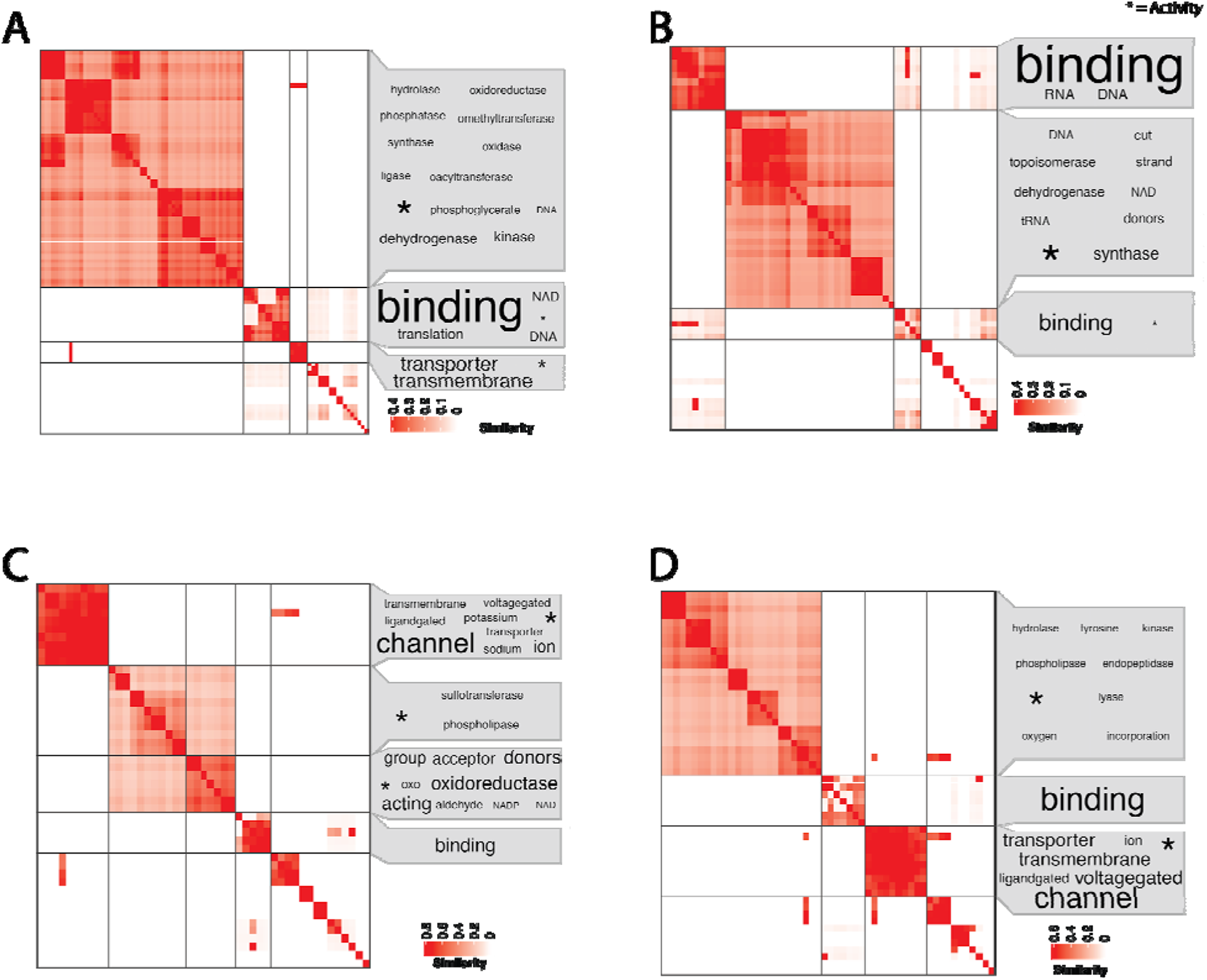
Clustered (binary cut) semantic similarity matrix of Molecular Function gene ontology terms for all genes in the 9 modules identified as the A) maternal mRNA complement (n=92 terms), the 11 modules of the B) first wave of the MZT (n=55 terms), the 10 modules of the C) second wave of the MZT transition (n=22 terms), the 7 modules of the D) “adult” expression (n=31 terms). Terms representing each cluster are shown in a word cloud, with size of term representing frequency of occurrence in the terms. The generic term, “Activity” (identified by * in each graphic), was put as a footnote for visual clarity.

### Expression of Developmental Biomarkers

The expression profiles of six developmental biomarker genes involved in the MZT and later development were assessed to test if our WGCNA-based timeline of the MZT was supported by these key developmental players. Several orthologs of MZT biomarkers were found using BLAST, including Cyclin-B (7 hits), Smaug (3 hits), Kaiso (90 hits), Sox2 (18 hits), Wnt8 (23 hits), and TBXT (15 hits). The expression profiles of these six genes, all of which showed significantly different expression patterns across life stages (Table S4A), indicate their specific temporal activity during early development (Fig 6). Firstly, Cyclin-B is highly expressed in unfertilized eggs up until cleavage. However, after this point its expression decreases monotonically until the planula stage (Fig. 6i). Next, both Smaug and Kaiso experience significant upregulation after fertilization and downregulation after the prawn chip stage (Fig. 6ii-iii). Then, during the early gastrula stage the zygotic transcription activator, Sox2, significantly peaks after which point it is rapidly downregulated (Fig. 6iv). Wnt8 appears to be lowly expressed in early on and is downregulated further after cleavage. However, it later significantly peaks in expression between the early gastrula and planula stages of development (Fig. 6v). Finally, TBXT is significantly upregulated starting after cleavage, leading to an expression peak that lasts until after the planula stage (Fig. 6vi).

**Figure 6.**
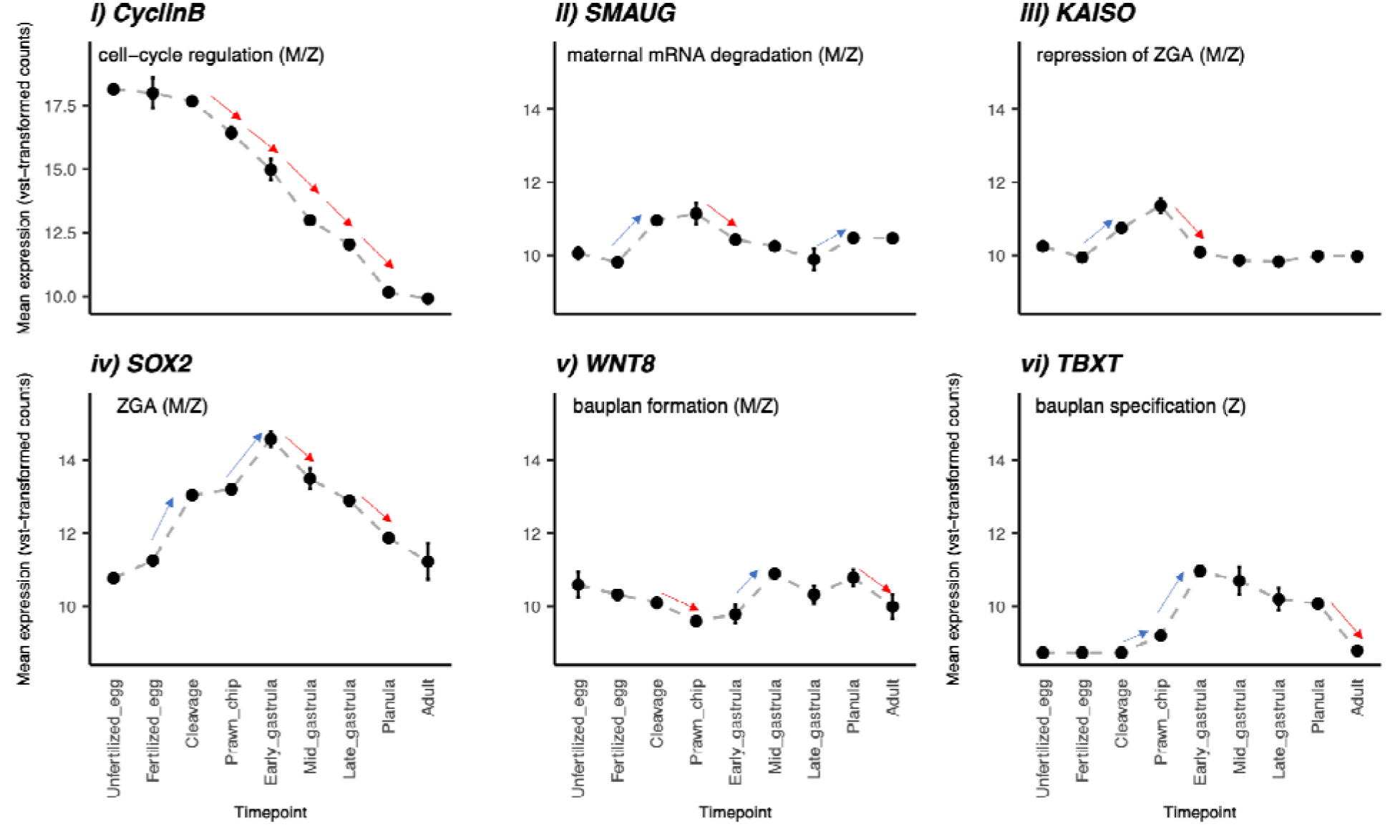
Expression of key biomarkers (M. capitata gene id) in the MZT: i) Cyclin-B (g71356), ii) Smaug (g4639), iii) Kaiso (g60350), iv) Sox2 (g53225), v) Wnt8 (g33149) and vi) Brachyury (TBXT; g68947). Points and error bars display mean±standard error of the mean. Red and blue arrows indicate transitions with significant (padj<0.05, log2FoldChange>|1|) down- and up-regulation, respectively. M=maternal expression, Z=zygotic expression, ZGA=Zygotic genome activation.

### Expression of Epigenetic Biomarkers

Targeted expression analysis of seven epigenetic biomarkers was also conducted to elucidate the epigenetic landscape necessary for the MZT. A Blast search of the *M. capitata* transcriptome reported 3 DNMT3a hits, 4 DNMT1 hits, 1 TET1 hit, 2 MBD2 hits, 1 MBD3 hit (shared with MBD2), 2 UHRF1 hits, and 84 BRG1 hits. The top hits of these seven genes all showed significantly different expression patterns across life stages (Table S7). Temporal expression of these epigenetic enzymes revealed three distinct patterns in expression (Fig 7). First, for all seven enzymes, expression levels changed very little between the samples representing unfertilized eggs and fertilized eggs. After fertilization, DNMT1, MBD2/3, and UHRF1 decrease in expression throughout development (Fig. 7ii, 7iv, 7v). For DNMT1, while expression appears to decrease overall throughout development, a small peak in expression occurs during late gastrulation. For MBD2/3, expression appears to plateau after the initial decrease following fertilization, while expression of UHRF1 appears to decrease monotonically throughout development (Fig. 7iv, 7v). In contrast to DNMT1, MBD2/3, and UHRF1, the expression of DNMT3A, TET1 and BRG1 increases significantly after fertilization and later plateaus. For TET1 and DNMT3A transcription appears to stabilize after cleavage, however BRG1 continues to be significantly upregulated until the prawn chip stage before plateauing.

**Figure 7.**
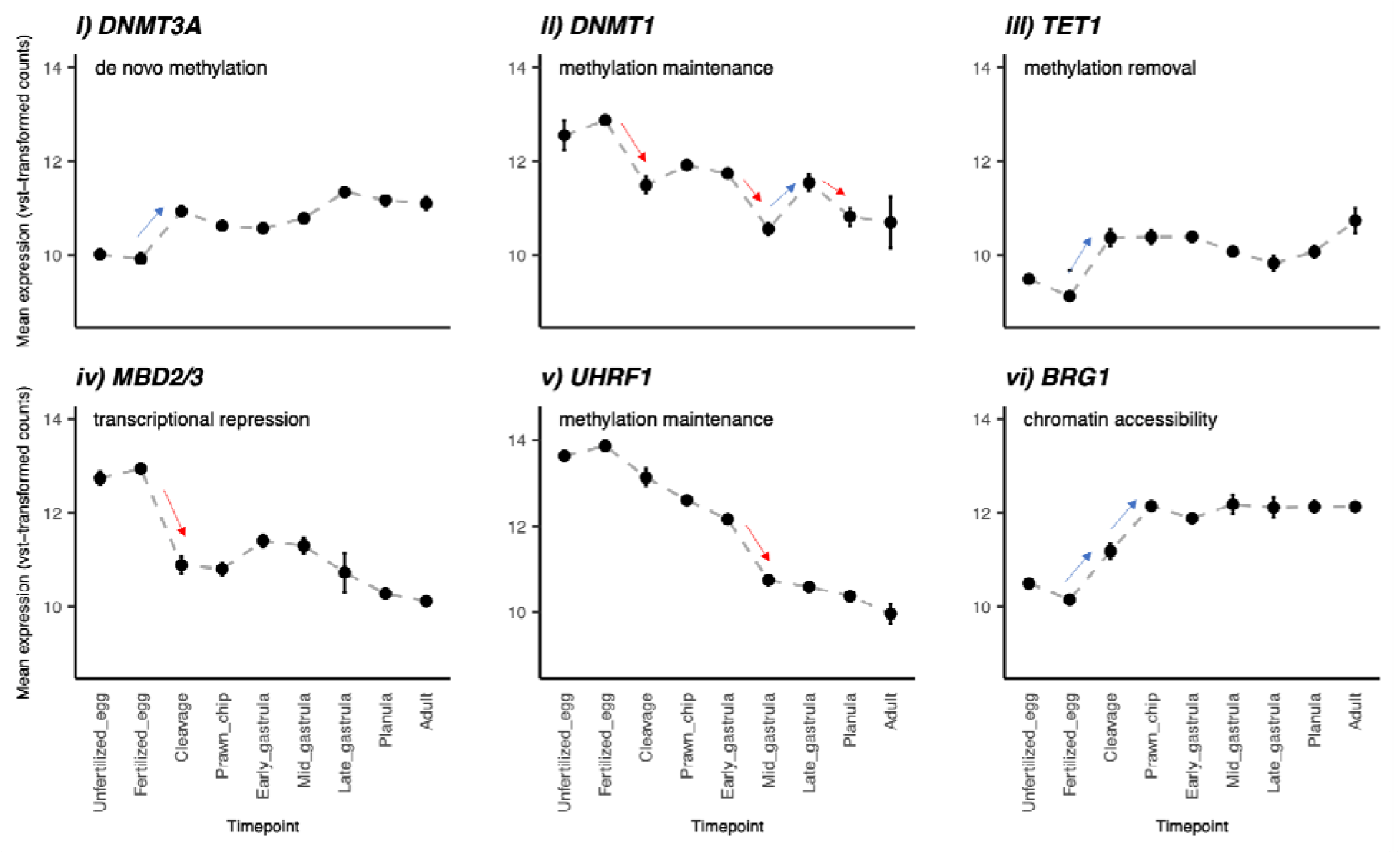
Expression of enzymes (M. capitata gene id) involved with i) DNMT3a (g25804), ii) DNMT1 (g53952) iii) TET (adi2mcaRNA27872_R0), iv) MBD2 and MBD (both g53132), v) UHRF1 (adi2mcaRNA19502_R1) and vi) BRG1 (g68733). Points and error bars display mean±standard error of the mean value of the VST normalized data. Red and blue arrows indicate transitions with significant (padj<0.05, log2FoldChange>|1|) down- and up-regulation, respectively.

## Discussion

In this study, developmental gene expression of the reef-building coral, *Montipora capitata* was examined to characterize the maternal mRNA complement and function in oocytes and the timeline of the maternal to zygotic transition (MZT). In sampling nine time points, with six time points occurring within thirty-six hours of fertilization, our dataset provides the highest temporal resolution to date that captures complex gene expression dynamics underlying the MZT. WGCNA facilitated the identification of the maternal mRNA complement, two waves of ZGA, and adult expression. These results, along with targeted expression analysis of six developmental biomarkers provides evidence that the MZT begins before cleavage (4 hpf) and continues through gastrulation. The expression profiles of seven epigenetic enzymes provide further support for the timing of the MZT and the epigenetic state necessary for ZGA in *M. capitata*. These findings provide an essential framework for future research on mechanisms of resilience in early developmental stages and the potential for parental carryover effects under increasing environmental stressors.

### Maternal Provisioning

The active maternal mRNA complement, here defined as the modules significantly positively correlated with either the unfertilized and fertilized egg samples, was enriched in biological processes primarily related to housekeeping functions (e.g. vacuolar transport [GO:0007034], spliceosomal snRNP assembly [GO:0000387], D-amino acid metabolic process [GO:0046416], and DNA replication [GO:0006260]). This is unsurprising as the primary function of maternal mRNA provisioning is to regulate early development until the molecular machinery of the zygote is able to take over [23]. Extensive work across animal phyla shows enrichment of housekeeping functions in eggs [26,74–76]. For example, in a study comparing the maternal mRNA complement of *Lymnaea stagnalis* with maternal transcriptomes across the animal kingdom, it was found that the 5-10% of maternal mRNAs conserved across phyla consisted primarily of functions such as nucleotide binding, protein degradation and activities associated with the cell cycle [26]. Indeed, in our analysis via WGCNA, we found support for significantly enriched GO terms (Table S3.) commonly associated with cell cycle, biosynthesis, transcription, signaling, and protein processing functions. This reinforces results from a prior GO analysis via enrichment of genes expressed in both female and male gametes with 100TPM threshold from these same three unfertilized egg samples [33].

High representation of mRNAs involved in DNA damage repair in the maternal complement (e.g., DNA damage checkpoint [GO:0000077], DNA repair [GO:0006281], recombinational repair [GO:0000725], interstrand cross-link repair [GO:0036297], cellular response to ionizing radiation [GO:0071479], and positive regulation of apoptotic process [GO:0043065]) likely prevents the proliferation of damaged cells during the rapid cell cycles occurring during the first few hours of development. Embryonic development is particularly sensitive to DNA damage, as the accumulation of damaged cells can quickly lead to embryonic death [77–79]. Embryos of broadcast spawning species, especially those of vertically transmitters that pass down symbionts in the eggs [80], are particularly at risk of DNA damage due to exposure to ultraviolet (UV) radiation [81–84]. UV radiation can damage DNA both indirectly, through the production of reactive oxygen species that can lead to the oxidation of nucleotides, and directly by dimerization of neighboring pyrimidines [85–87]. Prolonged exposure to UV-radiation during early development may necessitate maternal frontloading of DNA repair mechanisms in broadcast spawners [81–84]. For example, its been shown that anchovy eggs spawned under light and dark conditions have an innate capacity for UVB-induced DNA repair, but that diel cycles of DNA repair mechanisms (likely zygotically-derived) did not occur until approximately six hours post-fertilization [84]. Furthermore, Antarctic meroplankton species that spawn during the Austral summer have been shown to have a higher capacity for DNA damage repair than species that spawn during the winter [83]. As a tropical broadcast spawning species with vertical transmission of symbionts [67,88], *Montipora capitata* embryos are particularly susceptible to DNA damage incurred from UV radiation and from ROS production by their endosymbionts [80,89–91], which may partially explain the enrichment of DNA repair functions in the *M. capitata* maternally-provisioned oocyte transcriptome.

While the gene expression time course analyzed here provides insight into the maternal complement, we acknowledge that our analysis may not be comprehensive. The expression of maternal mRNAs in the embryo is primarily regulated through the modulation of the poly-A tail [23,39,41,92,93], with elongation of the poly-A tail leading to transcriptional activation and shortening of the poly-A tail leading to de-activation [23,39,41,92,93]. As our sequence library preparation (polyA enrichment) primarily excludes un-adenylated mRNAs, it is likely that some un-adenylated transcripts, potentially related to stress-response functions in the maternal mRNA complement, were not fully captured in these libraries. Absolute determination of maternal mRNAs (adenylated and un-adenylated) in the oocytes could be better served via ribosomal RNA depletion library preparation prior to sequencing, as suggested in [40].

### MZT Timeline

With the exception of non-parthenogenetic insects, in most metazoans, fertilization triggers egg activation and is associated with the elimination of approximately 30% of maternal transcripts [39,49,94,95]. The clearance of these maternal mRNAs marks the first event of the MZT [39,49]. In *M. capitata*, a wave of maternal transcript degradation, marking the beginning of the MZT, appears to be starting by 1-4 cell divisions (4 hpf). As sampling time increases beyond 1 hpf, the maternal complement (modules grey, coral1, lightslateblue, mediumpurple3, antiquewhite2, antiquewhite4, thistle, honeydew1, and midnightblue) shows a substantial reduction of correlation of eigengene expression with life stage (Fig. 3), supporting the downregulation of transcripts that are associated with unfertilized and fertilized eggs. This is further evidenced by the significant downregulation of maternally-provisioned Cyclin-B mRNAs starting by 4 hpf (Fig. 6i), and a peak in the expression of Smaug (Fig. 6ii) between cleavage (4 hpf) and prawn chip (9 hpf), signaling the de-adenylation of maternal transcripts during this period. After the clearance of maternal mRNAs between fertilization and cleavage, the clustering of life stages by module eigengene correlation (Fig. 3) supports two temporal peaks in expression, suggesting that in *M. capitata*, the MZT progresses in two waves. The first wave appears to occur between 4 and 14 hpf, as indicated by the clustering of eigengene-lifestage correlations of cleavage, prawn chip, and early gastrula samples, while the clustering of mid-gastrula, late gastrula, and planula samples support a second wave occurring between 22hpf and 9dpf. While we hypothesize the ZGA is ongoing until settlement, as indicated by the unique expression pattern in the adult transcriptome, the transfer of developmental control to the zygote, marking the conclusion of the MZT, likely occurs much earlier. However, the discrimination of transcripts as “maternal” or “zygotic” in origin can be complicated because the activation of existing mRNAs by polyadenylation cannot be distinguished from the activation of newly-transcribed mRNAs, as discussed above. As such, we cannot distinguish the absolute end of the MZT, which is marked by the transition of developmental control from maternal to zygotic molecular machinery [39]. In the future, the termination of the MZT could be more fully captured through the characterization of Single Nucleotide Polymorphisms (SNPs) in the maternal and paternal transcriptomes, as described in [96], zygotically-derived transcripts can be identified by the presence of paternal SNPs. For corals, this would best be accomplished through making crosses between eggs and sperm from known parental colonies.

Taken in concert, the expression profiles of key developmental enzymes provide a temporal progression illustrating the molecular mechanisms underlying zygotic genome activation in *M. capitata*. Peaks in the expression of Cyclin-B, Smaug, Kaiso, and Sox2 (Fig 3i-3iv) indicate that maternal mRNA clearance and ZGA during the first wave of the MZT in *M. capitata* is mediated by maternal mRNAs. Indeed, the polyadenylation of maternal mRNAs after fertilization have been implicated in both maternal mRNA degradation [45,46,51] and ZGA [50,53,97] in bilaterians. The expression profiles of these enzymes during *M. capitata* and sea anemone (*Nematostella vectensis*) MZT [98] supports that maternal regulation of the first MZT wave is a conserved feature from basal metazoan development. Further, the peak in maternal transcriptional regulator Kaiso in the prawn chip samples (9 hpf), followed by the peak in maternal transcription factor Sox2 in the early gastrula samples (14 hpf) suggests that ZGA is repressed until early gastrulation in *M. capitata*. This is further supported by a substantial increase in the correlation of eigengene expression with time during and after the early gastrula stage in module clusters 5, 8, and 9 (Fig. 3 Y axis), and a gradual shift in dominance of the enrichment of RNA and protein processing-related biological processes in the cleavage stage (Fig. S2C, Table S2C) to terms related to transcription in the early gastrula stage (Fig. S2E, Table S2E).

Several factors including nucleocytoplasmic ratio, chromatin accessibility, cell cycle destabilization, and maternal mRNA degradation have been implicated in the initiation of zygotic transcription [39]. In *N. vectensis*, yeast, and the African clawed frog cell cycle destabilization appears to facilitate the activation of zygotic transcription [98–101]. In *Nematostella vectensis*, this is shown by a decrease in Cyclin-B between 2 and 7 hpf followed by ZGA between 7 and 12 hpf. In *M. capitata,* Cyclin-B also decreases after cleavage (4 hpf), showing cell cycle destabilization. While cell cycle destabilization likely plays a role in ZGA in *M. capitata*, increasing chromatin accessibility (see “Epigenetic state for ZGA”) and the weakening of maternal transcriptional repression through the de-adenylation of maternal mRNAs, indicated by a peak in RNA binding protein Smaug between 4 and 9 hpf, likely also contribute to ZGA.

While the first wave of the MZT appears to be regulated through maternal mRNAs, a shift in positive eigengene correlations from modules in cluster 4 to modules in clusters 8 and 9 between early and mid-gastrulation (Fig 3) shows the upregulation of genes that were either previously lowly expressed, or completely unexpressed between 14 and 22 hpf. A significant decrease in the expression of Sox2 during this transition indicates that zygotic, rather than maternal, transcription factors may be implicated, supporting the identification of this temporal peak in gene expression as the second wave of ZGA. While expression of Wnt8 and TBXT peaks during this second wave is expected due to their roles in bauplan specification during gastrulation [98,102,103], their continual upregulation despite the downregulation of Sox2 suggests that they may be zygotically transcribed in *M. capitata,* as shown in *Xenopus* and in sea urchins. [53,54,97]. Besides bauplan specification, similarly to [98] ion/peptide transport and cell signaling-related processes are upregulated during the second wave (Fig 4C). Interestingly, we also see upregulation of GO terms related to response to environmental stress (i.e. xenobiotic transport (GO:0042908) and response to oxidative stress (GO:0006979)), as well as terms related to symbiosis (i.e. reduction of food intake in response to dietary excess (GO:0002023), response to glucose (GO:0009749), multicellular organism growth (GO:0035264), glycerolipid biosynthetic process (GO:0045017), carbohydrate biosynthetic process (GO:0016051) and glycolipid biosynthetic process (GO:0009247)), in the late gastrula (Table S2G) and planula stages (Table S2H), suggesting that by late gastrulation (60 hpf) *M. capitata* embryos are able to sense the environment and tune transcription in response to environmental stimuli. The combined loss of maternal defenses at the start of the MZT and low homeostatic capacity has been suggested to contribute to embryo vulnerability in fish, sea urchins, and other marine broadcast spawners [6,7,18–22]. As such, the period ranging from the start of maternal mRNA clearance (≤4 hpf) to transcriptional plasticity in response to stimuli (≤60 hpf) may represent the most sensitive stages and thus present a critical bottleneck in *M. capitata* early development.

While the timing and scale of maternal transcript degradation and ZGA during the MZT is shown to vary between species [40], the timing of MZT described here is remarkably similar to other anthozoans with similar developmental courses. For example, gene expression time courses during *Acropora digitifera* [34], *Acropora millepora* [35], and *Nematostella vectensis* [98] development suggest the onset of ZGA occurs within the first 24 hours of development. In *A. digitifera*, clustering of the expression patterns in early life stages mirrored the clustering of life stages here [34]. In *A. digitifera*, the prawn chip (9 hpf) and early gastrula (14 hpf) stages grouped together, and late gastrula (48 hpf), planula (96 hpf), and adult stages grouped together [34]. Likewise, a shift in gene expression between cleavage (4 hpf) and gastrulation (22 hpf) in *A. millepora* supports the onset of ZGA during the early gastrula stage [35]. The sea anemone, *N. vectensis*, also shows similar gene expression dynamics during the MZT to *M. capitata*, with maternal degradation in *N. vectensis*, potentially occurring between cleavage (2 hpf) and prawn chip (7 hpf), and ZGA beginning by the onset of gastrulation (12 hpf; [98]. While the genera *Acropora* and *Monitpora* belong to the same family of reef-building corals (Acroporidae), *N. vectensis* belongs to different sub-class in the anthozoan lineage. However, the four species are all broadcast spawning anthozoans with similar developmental timelines and embryonic courses (i.e. form a prawn chip; [34,68,104,105], suggesting that the timing of the MZT, and potentially the molecular mechanisms underlying MZT regulation, may be evolutionarily conserved within cnidarians. If so, this would have important implications for research on carryover effects and transgenerational plasticity, which depends heavily on the rapid accumulation of knowledge surrounding gene expression regulation and epigenetic state during reef-building coral development [56,106]. Much of this knowledge has already been generated in the model cnidarian, *N. vectensis* [98,107], and may provide a strong baseline for such studies.

### Epigenetic state for ZGA

Epigenetic programming during the MZT has a key role in the repression and activation of zygotic transcription [40,55], and additionally presents a potential mechanism for heritable, non-genetic phenotypic plasticity allowing for the rapid acclimatization of reef-building corals to rapid environmental change [56,106]. However, epigenetic programming during the MZT has primarily been described in mammals and non-mammalian vertebrates and likely differs greatly from epigenetic programming during the development of reef-building corals and other invertebrates [108–111]. In mammals, parental epigenetic marks are erased twice during the MZT, and are replaced with marks that are necessary for embryo viability [112,113], making epigenetic inheritance rare in mammals [108]. However, nonmammalian vertebrates appear to inherit the paternal methylome, reprogramming the maternal methylome to match before ZGA begins [114]. Epigenetic programming during the MZT is only starting to be understood in invertebrates, as historic invertebrate models (e.g., *Drosophila melanogaster and Caenorhabditis elegans*) lack cytosine DNA methylation almost entirely [114], and the invertebrate species studies thus far appear to lack epigenetic reprogramming entirely (e.g., sponges, honey bees, ctenophores, sea urchins, and sea squirts; [108,109,111,115]. Indeed, a recent work on reef-building corals shows that epigenetic marks may be transferred between generations, and suggests that the paternal and maternal methylomes appear to contribute equally to the methylome of the offspring [110].

Generally in vertebrates and mammals in particular, chromatin structure is relatively open prior to ZGA, and decreases in accessibility during the MZT as DNA methylation increases [40,116,117]. While chromatin structure condenses over time, local DNA accessibility remains open during ZGA to facilitate the activation of specific genes [40,116,117]. However, this appears to not be the case for invertebrates. For example, the bivalves *Crassostrea gigas* and *Patinopecten yessoensis*, are shown to have heavy methylation in the oocytes, which peaks during the blastula/gastrula stage before declining and remaining at stable levels for the remainder of development [118,119]. In other invertebrates, such as honey bees and sponges, DNA methylation levels tend to be stable throughout early development [109,111,115]. While methylation levels were not measured here, significant changes in the expression enzymes linked to methylation activity may hint to the epigenetic landscape necessary for ZGA in *M. capitata*.

The expression patterns of key the epigenetic regulators assessed here provide the first data characterizing the dynamics of genes involved in epigenetic programming underlying the MZT in *M. capitata* and further sets the stage for future research on the mechanisms of epigenetic inheritance in reef-building corals. Here, significant changes in the expression of six out of seven of these genes between the samples representing fertilized eggs (1 hpf) and cleaving embryos (4 hpf) correlates with the start of the MZT. DNMT3a and TET1 responsible for de novo methylation and methylation removal, respectively, are two of these six enzymes that change between 1to 4 hpf and then maintain stable expression for the remainder of development, suggesting that the embryo methylome programming occurs during this short window. Completion of methylome programming may be necessary for the start of the MZT, as it has been shown in zebrafish that harmonization of the paternal and maternal methylomes is necessary for the successful progression of ZGA [120]. A sharp decrease in transcriptional repression through MBD2/3 by the cleavage stage may signal the completion of methylome harmonization. Accordingly, research on the mechanisms underlying the transfer of epigenetic marks from parent to offspring and harmonization of the embryonic methylome in reef-building corals may benefit from frequent sampling and quantification of methylation between fertilization (0 hpf) and cleavage (4 hpf).

Several mechanisms of epigenetic modification appear to set the stage for ZGA after embryonic methylome programming in *M. capitata*. Firstly, passive (de)-methylation is likely to be the primary regulator of DNA methylation during the MZT due to rapid cell division. While expression of DNMT3a (*de novo* methylation) and TET1 (methylation removal) stabilizes after cleavage, DNMT1 (methylation maintenance) shows significant expression fluctuations during the MZT. After initial down-regulation of DNMT1 prior to cleavage, methylation maintenance activity appears to remain stable through the first wave of the MZT. However, it is significantly downregulated again at the start of the second wave of the MZT (during mid-gastrulation), prior to returning to previous expression levels. This depression of methylation maintenance provides support to the hypothesis that increasing local chromatin accessibility facilitates the activation of specific loci [40,118,121]. Increasing local chromatin accessibility is further evidenced by upregulation of BRG1 from pre-cleavage (1 hpf) until the prawn chip stage (9 hpf), which may additionally facilitate the start ZGA during early gastrulation. Other mechanisms of epigenetic modification with putative roles in the MZT [49], including histone arginine and lysine methylation (GO:0034969, GO:0034968), histone acetylation (GO:0016573), and protein ADP-ribosylation (GO:0006471) were enriched in the maternal complement (Table S2) and during the first wave of ZGA (Table S3), supporting multiple conserved mechanisms of epigenetic regulation of the MZT between model organisms and *M. capitata*.

## Conclusions

As anthropogenic climate change continues to alter the marine environment at an accelerating and unprecedented rate, it is more important than ever to understand the mechanisms underlying the resilience and sensitivity of vulnerable ontogenetic stages. Future research evaluating potential avenues of rapid acclimatization in reef-building corals, including parental effects, carryover effects, and cross-generational plasticity, relies on the generation of baseline knowledge on parental molecular provisioning, the epigenetic landscape during early development, and temporal gene expression dynamics during the MZT [56]. Here, our characterization of maternal mRNA provisioning and the maternal-to-zygotic transition in *Montipora capitata* sets the stage for these future studies and additionally identifies cleavage (≤4 hpf) through late gastrulation (≤60 hpf) as a potential critical window during *Montipora capitata* development.

## Methods

### Study System

Endemic to the Hawaiian archipelago, *Montipora capitata* is a dominant and ecologically important inhabitant of lagoons and fringing reefs [67,122–124]. Studies have shown that *M. capitata* is more tolerant to environmental stressors compared to other reef-building corals due to its perforate skeleton and deep tissues [125,126]. However, the earliest life stages of this species may be more susceptible to climate change-related and local anthropogenic stressors than adults. Like many other marine invertebrates, *M. capitata* has a complex early life history. It is a hermaphroditic broadcast spawner with vertical symbiont transmission [67,88]. The early life history of *Montipora spp.*, from fertilization through larval body formation, has been well-documented by Okubo and others [68].

### Specimen collection

*M. capitata* egg-sperm bundles were collected (Hawai□i Department of Land & Natural Resources Special Activity Permit 2018-50) from the reef adjacent to the Hawai□i Institute of Marine Biology (HIMB) Kāne□ohe Bay, Hawai□i (21°25′58.0″N 157°47′24.9″W) during their release on June 13, 2018. *M. capitata* adult fragments were collected from the reef adjacent to HIMB and Reef 11.13 (1°27′07.7″N 157°47′40.3″W) and acclimated in ambient conditions (27 °C, ∼480 µatm) mesocosm tanks for two weeks prior to snap freezing on September 22nd, 2018.

### Experimental set-up

Immediately after collection, 300 uL of egg and sperm bundles were snap frozen and stored at − 80 °C while the rest of the bundles were placed in conical chambers to break apart and hydrate for 10 minutes. 300 uL of eggs were separated from the sperm and rinsed 3 times with 0.2 uM filtered seawater then snap-frozen and stored at −80 °C. Fertilization and early development (0 h − ∼16 hfp) took place in 3 replicated 1.6L flow through conical chambers and further development (∼ 16 hpf − 9 d) took place in small (6 x 6 cm) flow through bins within larger 74L flow through tanks, both under ambient temperature ∼26.8°C. Water flow to each conical was controlled with ½ GPH Pressure Compensating Drippers and held at a maximum potential flow rate of 7.57 liters per hour. For a 1.6 L conical, the turnover rate was once every ∼50 minutes and for a 74L tank, the turnover rate was once every ∼ 23 hours. The average light intensity of the experimental setup was measured using a handheld PAR sensor (Underwater Quantum Flux Apogee instruments - Model MQ-510; accuracy = ±4%) and was 115.8 ± 14.7 µmol m^-2^ s^-1^, n=6 measurements. Temperature, salinity, and pH were measured three times daily using a handheld digital thermometer (Fisherbrand Traceable Platinum Ultra-Accurate Digital Thermometer, accuracy = ±0.05 °C, resolution = 0.001°) and a portable multiparameter meter (Thermo Scientific Orion Star A-series A325; accuracy = ±0.2 mV, 0.5% of PSU reading, resolution = 0.1 mV, 0.01 PSU) with pH and conductivity probes (Mettler Toledo InLab Expert Pro pH probe #51343101; Orion DuraProbe 4-Electrode Conductivity Cell Model 013010MD), respectively. Temperature and pH probes were calibrated using Tris (Dickson Laboratory Tris Batch 27, Bottles 70, 75, 167, 245, and 277) standard calibrations. Throughout the experiment, the temperature, salinity, and pH of tanks and conicals (mean±SEM) was 26.99±0.075°C, 34.24±0.017 psu, and −52.84±0.348 mV, respectively.

### Sample collection and preservation

Samples were collected at nine time points. Sampling times (Fig. 1) represent visually-distinct developmental stages including unfertilized eggs (immediately after bundle breakup; n=3) and multiple stages at various hours post fertilization (hpf), fertilized egg (1 hpf; n=2), cleavage (4 hpf; n=3), prawn chip (9 hpf; n=3), early gastrula (14 hpfg; n=3), mid-gastrula (22 hpf; n=2), late gastrula (1.5 days post fertilization; n=2), and planula (9 days post fertilization; n=3). Molecular samples, 300 µL/replicate of gametes, embryos, and larvae, or whole adult fragments (∼5×7 cm; n=3), were snap-frozen in liquid nitrogen and stored at −80°C until RNA extraction.

### RNA extraction, sequencing, and read processing

Developmental samples were digested at 55°C in 300 µL DNA/RNA Shield for two to three-and-a-half hours and centrifuged at 2200 rcf for one minute to separate the remaining solids. A small clipping of each adult sample was vortexed in 2 mL 0.5 mm glass bead tubes (Fisher Scientific Catalog No. 15-340-152) with 1 mL of DNA/RNA Shield for two minutes at high speed. Total RNA was extracted from each supernatant with the Zymo Quick-DNA/RNA™ Miniprep Plus Kit (Zymo Research, Irvine, CA, USA) following the manufacturer’s protocol for tissue samples. RNA was quantified with a ThermoFisher Qubit Fluorometer and quality was measured with an Agilent TapeStation 4200 System. Total RNA samples were then sent to Genewiz (South Plainfield, New Jersey, USA) for library preparation and sequencing. cDNA libraries were constructed following the TruSeq Stranded mRNA Sample Preparation Protocol (Illumina) using polyA enrichment and were sequenced on a HiSeq instrument at Genewiz targeting 15 million reads per sample.

Read quality was assessed with FastQC (v0.11.8) and compiled with MultiQC [127,128]. Following quality assessment, reads were trimmed to remove adapters and low-quality reads using FastP [129]. Sequences were filtered for quality applying a five base pair sliding window to remove reads with an average quality score of 20 or less. The sequences retained had quality scores greater than or equal to 20 in at least 90% of bases and a sequence length greater than or equal to 100 bases. Read quality was re-assessed with FastQC (v0.11.8) and then aligned and assembled using HISAT2 in the stranded paired-end mode [130] in combination with StringTie in the stranded setting (v2.1; [131]. Following assembly, mapped GFFs were merged for assessment of precision and accuracy and compared to the *M. capitata* reference assembly GFF using GFFcompare (v0.11.5; [132]. A gene count matrix was then generated from the GFFs using the StringTie *prepDE* python script [131].

### Global gene expression analysis

All gene expression analyses were performed in RStudio (v1.3.959), using R version 4.0.2 [133]. First, genes with low overall expression were filtered using Genefilter’s (v1.70.0) *pOverA* filter function [134]. Given that the smallest number of replicates was two, genes with fewer than 10 counts in at least 2 out of 24 samples (*pOverA* 0.083, 10) were excluded from further analysis. Counts were then normalized with DESeq2’s (v1.28.1) variance stabilizing transformation (vst; [135] after confirming that all size factors were less than 4. To visualize experiment-wide patterns in gene expression, a principal coordinates analysis (PCA) based on sample-to-sample distances was performed on the vst-transformed gene counts using the DESeq2 *plotPCA* function (Fig. 2).

Transformed counts were used for a Weighted Gene Co-expression Network Analysis (WGCNA; [136]. WGCNA analysis is a data reduction approach that assigns genes with similar expression patterns to co-expression groups called modules. Using this technique allowed us to identify groups of genes with similar expression profiles across developmental time. All samples were used for WGCNA after visually checking for outliers in an unrooted hierarchical tree built with the R stats *hclust* “average” function [133]. In order to construct a topological overlap matrix similarity network to assess gene expression adjacency, the WGCNA *pickSoftThreshold* function was used to explore soft threshold values from 1 through 30. A soft thresholding power of 23 (scale-free topology fit index of 0.85) was chosen and used to construct a topological overlap matrix similarity network using signed adjacency. Modules were identified from this topological overlap matrix similarity network using the WGCNA package *dynamicTreeCut* function with the settings deepSplit 2, and minimum module size 30. Modules with greater than 85% eigengene similarity were merged and the resulting finalized 34 modules were used for expression plotting, module-trait correlation, and gene ontology enrichment analysis.

Clustering of WGCNA expression modules by eigengene dissimilarity using the *hclust* “average” method identified 9 clusters of modules. Expression profile plots were generated for each cluster by plotting the mean eigengene expression value of each of the replicate samples per time point for each module (Fig. S1). Additionally, module-trait correlation was assessed by calculating gene significance (the correlation between the gene and the time point) and module membership (the correlation of the module eigengene and the gene expression profile; [136]. Correlation values were plotted as a heatmap using the *complexHeatmap* package [137]. Modules with a p-value less than or equal to 0.05 were considered significantly correlated with a timepoint (Fig. 3). Finally, the distinct expression phases in the MZT were identified by clustering life stages by their module-trait correlation using the *hclust* “average” method to build an unrooted hierarchical tree.

### Gene ontology mapping and enrichment analysis

Comprehensive gene ontology (GO) annotation of the reference genome was undertaken for subsequent functional enrichment analysis using InterProScan (v.5.46-81.0), Blast2GO (v5.2), and UniProt [71–73]. First, homologous protein sequences were identified using the DIAMOND (v2.0.0) *blastx* program in “more sensitive” mode to map predicted cDNA sequences against the NCBI non-redundant (nr) protein database (downloaded on August 6, 2020) using an e-value cut-off of 1e-05 and a block size of 20 [70]. Concurrently, InterProScan’s *iprlookup* function was run to map GO terms from the InterPro database (accessed on August 24, 2020) to the *M. capitata* predicted protein sequences [69,71]. Next, the XML output files from DIAMOND and InterProScan were both loaded into Blast2GO for compilation and further mapping using the obo database (updated August 11, 2020; [72]). Additional GO terms were extracted from the UniProtKB database by searching for protein identifiers obtained via DIAMOND using UniProt’s “Retrieve/ID mapping tool” [73]. Finally, mapping results from Blast2GO and UniProt were compiled in RStudio (v1.3.959), using R (v-4.0.2; [133]).

Gene ontology enrichment analysis was performed to characterize the functions provided by the maternal mRNA complement and the changes in the embryo’s functional toolkit during the MZT. Modules that were significantly correlated (p<0.05) with the maternal and adult complements and each phase of the MZT, as well as each life stage, were identified. The genes comprising each module within a group of interest were compiled and subsetted from the counts matrix for functional enrichment analysis. The R package *Goseq* (v1.40.0) was used to perform GO term enrichment analysis, therefore taking gene length bias into account [138] when determining the functional profiles of each group. Subsequently, the R package *simplifyEnrichment* (v0.99.4) was used to visualize the enriched biological process and molecular function terms for each group [139]. With this package, a semantic similarity matrix for a set of GO terms is generated based on GO tree topology, which is then used to cluster genes with ‘binary cut’ [139]. The functional characteristics of each cluster is then displayed with a word cloud. Accordingly, we used the *simplifyEnrichment* with the ‘Relevance’ measure of similarity to calculate semantic similarity and display the dominant biological processes and molecular functions enriched in the maternal and adult complements, each phase of the MZT, and each life [139,140].

### Expression of Key Developmental and Epigenetic Biomarkers

While WGCNA provides a global view of gene expression dynamics during the MZT, examining enzymatic expression of key MZT biomarkers provides a more detailed view of this process. In light of this, we examined the expression of enzymes linked to maternal transcript degradation (Cyclin-B, and Smaug), suppression of zygotic transcription (Kaiso), and zygotic genome activation (Sox2, Wnt8, and TBXT), as well as the expression of enzymes that provide transcriptional regulation capacity during the MZT (DNMT1, DNMT3A, TET1, MBD2, MBD3, UHRF1, and BRG1). A local alignment search of the *M. capitata* transcriptome was used to find the enzymes of interest for expression analysis [69,73]. Anthozoan (as available) and model organism protein sequences for selected enzymes were identified from the NCBI protein database for query. Sequence length, organism, quality, and date modified were considered in cases for which there were multiple available sequences. *Blastx* was used to identify the enzymes of interest in the *M. capitata* reference transcriptome (e-value < 10−5, max_target_seqs=100) using a Blast database created from the predicted cDNA sequences [69]. The hits table was filtered for duplicates in R. This table was used to subset the vst-normalized gene counts matrix, leaving only hits to the enzymes of interest. The top ten hits for each enzyme are reported in Table S8. The top hits for each enzyme generally exhibited much higher bitscores and lower e-values compared to subsequent hits. However, in some cases, the blast results and expression profiles for the top 2-3 hits were nearly identical, even though the genes were located on different scaffolds in the genome. Due to the robust alignment of the top hits to the *M. capitata* genome, and the similar expression profiles of comparable hits, vst-transformed expression was plotted only for each enzyme’s top hit. In plotting the expression of these developmental and epigenetic biomarkers, we gain further support on the timeline of and the epigenetic state necessary for the MZT.

The expression profiles of the top developmental and epigenetic enzymes are supported with differential expression analysis. To test for changes in expression between subsequent timepoints, a DESeq2 dataframe was first constructed using all genes that passed the *pOverA* (*pOverA* 0.083, 10) filter described above [134,135]. Differential expression analysis was then performed on filtered counts using the Wald model to estimate pairwise differences in gene expression between subsequent life stages [135]. Finally, significant changes in the expression of the developmental and epigenetic biomarkers were extracted from the DESeq2 results, by filtering for the top hits for each enzyme and applying a cut-off of >|1| logfoldchange and padj<0.05 (Table S8). These results indicate when transcriptional regulation occurs for these key enzymes, providing further support for their activity during the early development of *M. capitata*.

## Declarations

### Ethics approval and consent to participate

Not applicable

### Consent for publication

Not applicable

### Availability of data and materials

All raw data generated for this project has been deposited in the NCBI repository and can be accessed with Bioproject accession ID, <u>PRJNA616341</u>. Additionally, all code used for transcriptomic analysis is publicly accessible from the GitHub repository, <u>Mcapitata_Developmental_Gene_Expression_Timeseries</u>, which will be submitted for a DOI on Zenodo upon acceptance for publication.

### Competing interests

The authors declare that they have no competing interests.

## Funding

This work was funded by BSF grant 2016321 to HMP and TM. This work was also partially supported by the USDA National Institute of Food and Agriculture, Hatch Formula project accession number 1017848.

### Authors’ contributions

HMP and TM designed and provided financial support for the study; ES MN, VS, MS, and HMP conducted the field work and collected the samples; EC conducted the molecular work; EC and HMP analyzed the data; EC, HMP, and ES wrote the manuscript, all authors contributed to the final manuscript.

## Acknowledgements

We dedicate this work to our colleague and friend Dr. Diane Adams. We would like to thank the faculty and staff of the Hawai_LJ_i Institute of Marine Biology and the University of Rhode Island Computing.

## Notes

https://github.com/echille/Mcapitata_Developmental_Gene_Expression_Timeseries

https://www.ncbi.nlm.nih.gov/bioproject/?term=PRJNA616341

### Competing Interest Statement

The authors have declared no competing interest.

