## Supplementary Material for "Developmental series of gene expression clarifies maternal mRNA provisioning and maternal-to-zygotic transition in the reef-building coral *Montipora capitata*"

**Fig. S1.** Boxplot and overlaid points of mean eigengene expression value of each of the replicate samples per time point (n=3, except mid-gastrula and late-gastrula where n=2) for each WGCNA module cluster (A-I).

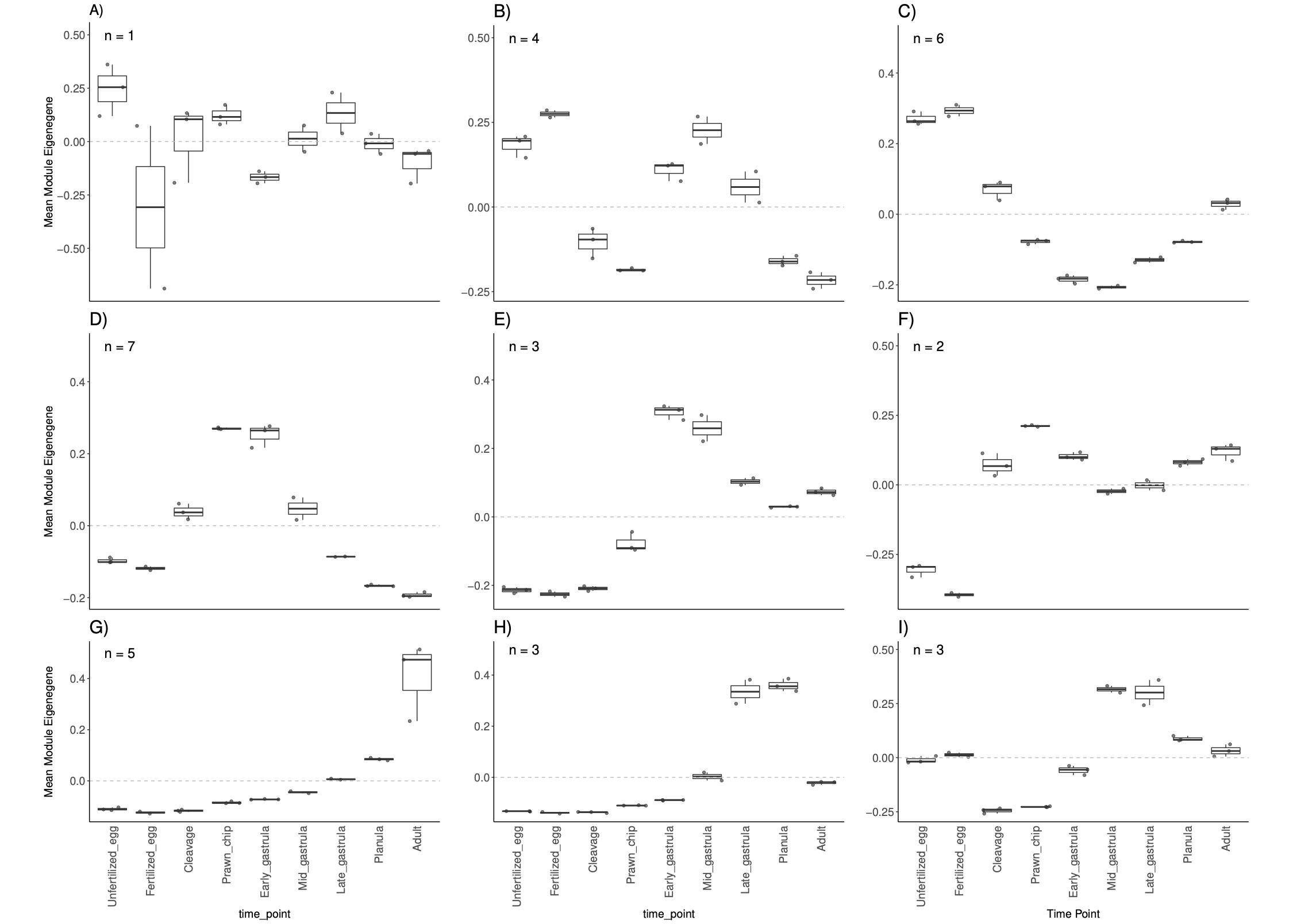

**Table S1.** Results from functional annotation of the genome using DIAMOND, InterProScan, Blast2GO, and Uniprot: [200824_Mcap_Blast_GO_KO.tsv](https://github.com/echille/Mcapitata_Developmental_Gene_Expression_Timeseries/blob/master/0-BLAST-GO-KO/Output/200824_Mcap_Blast_GO_KO.tsv)

**Table S2.** Significant biological process, molecular function, and cellular component gene ontology terms for each life stage. Results are sorted by time point and then by p-value of over-represented terms: [GO.05.allDev.csv](https://github.com/echille/Mcapitata_Developmental_Gene_Expression_Timeseries/blob/master/2a-WGCNA/Output/GO.05.allDev.csv)

**Table S3.** Significant biology process and molecular function gene ontology terms for the maternal mRNA complement. Results are sorted by time point, ontology, and p-value of over-represented terms: [GO.05.Mat.csv](https://github.com/echille/Mcapitata_Developmental_Gene_Expression_Timeseries/blob/master/2a-WGCNA/Output/GO.05.Mat.csv)

**Table S4**. Significant biology process and molecular function gene ontology terms for the first wave of the MZT. Results are sorted by time point, ontology, and p-value of over-represented terms: [GO.05.ZGA1.csv](https://github.com/echille/Mcapitata_Developmental_Gene_Expression_Timeseries/blob/master/2a-WGCNA/Output/GO.05.ZGA1.csv)

**Table S5**. Significant biology process and molecular function gene ontology terms for the second wave of the MZT. Results are sorted by time point, ontology, and p-value of over-represented terms: [GO.05.ZGA2.csv](https://github.com/echille/Mcapitata_Developmental_Gene_Expression_Timeseries/blob/master/2a-WGCNA/Output/GO.05.ZGA2.csv)

**Table S6**. Significant biology process and molecular function gene ontology terms for the second wave of the adult. Results are sorted by time point, ontology, and p-value of over-represented terms: [GO.05.Adult.csv](https://github.com/echille/Mcapitata_Developmental_Gene_Expression_Timeseries/blob/master/2a-WGCNA/Output/GO.05.Adult.csv)

**Table S7.** Differential expression analysis results from the 12 biomarkers genes (Figures 6 (A) and 7 (B)) with significant (padj<0.05, log2FoldChange>|1|) up- and down-regulation. Results are sorted by order by Figure 6 (A) and 7 (B), followed by life stage: [all_biomarker_DEGs.csv](https://github.com/echille/Mcapitata_Developmental_Gene_Expression_Timeseries/blob/master/2b-Epi-MZT-Enzyme-Expression/Output/all_biomarker_DEGs.csv)

**Table S8.** Top 10 hits to developmental biomarkers (A) and methylation-related enzymes (B) in the *M. capitata* transcriptome. Results are sorted by order in Figures 6 (A) and 7 (B) followed by bitscore. Only the first entry for each enzyme was plotted to examine expression profiles: [Epi_MTZ_biomarker_hits_Top10.csv](https://github.com/echille/Mcapitata_Developmental_Gene_Expression_Timeseries/blob/master/2b-Epi-MZT-Enzyme-Expression/Output/Epi_MTZ_biomarker_hits_Top10.csv)
